# EZH2 inhibition stimulates viral mimicry in resting splenic B cells

**DOI:** 10.1101/2023.05.08.539798

**Authors:** Seung J. Kim, Patti K. Kiser, Rodney P. DeKoter, Frederick A. Dick

## Abstract

In mammalian cells expression of repetitive genomic sequences is repressed by heterochromatin, underscoring the potential threat of repeat expression to cellular homeostasis. However, the specific consequences of ectopic repeat expression remains unclear. Here we demonstrate that EZH2 inhibitors stimulate repeat misexpression and cell death in resting splenic B cells. We show that B cells are uniquely sensitive to these agents because of high levels of H3K27me3 at repeats and correspondingly low DNA methylation. We generated a pattern recognition receptor loss-of-function mouse model called RIC with mutations in *Rigi*, *Ifih1* (MDA5), and *Cgas* to specifically block the consequences of repeat misexpression. In both WT and RIC mutant B cells, EZH2 inhibition caused focused loss of H3K27me3 at repetitive elements and upregulated their expression. However, expression of inflammatory chemokines and cell death were interrupted by the RIC mutations. Furthermore, the chemokine expression patterns induced by EZH2 inhibitors resemble the B cell response to Epstein-Barr virus infection. This study demonstrates a viral mimicry effect induced by pharmacological activation of repeat expression that induces inflammation and B cell death.

## Introduction

Enhancer of zeste homolog 2 (EZH2) is the catalytic subunit of the Polycomb Repressive Complex 2 (PRC2) that deposits di-and tri-methylation of histone 3 at lysine 27 (H3K27me2/3) ^1–3^. These are repressive histone modifications that cooperate with histone 2a lysine 119 ubiquitination (H2AK119ub) to silence transcription and compact chromatin ^4–11^. The latter modification is catalyzed by the Polycomb Repressive Complex 1 (PRC1), and the two PRCs were originally described in *Drosophila* as repressors of homeotic genes that dictate segmentation along the anterior-posterior axis ^12^. In mammals, the catalytic activity of EZH2 in PRC2 facilitates roles in cell fate determination ^13^, stem cell renewal^14^, and tumourigenesis ^15^. For example, EZH2 is required for B cell differentiation in the bone marrow during hematopoiesis ^16^, and germinal centre (GC) formation ^17^. Furthermore, EZH2 overexpression and gain-of-function mutations have been identified in different cancer types and are prominent in B cell lymphomas ^18–23^. Mechanistically, somatic mutations at EZH2^Y641^ render it dominantly active, increasing H3K27me3 globally, and repressing cell cycle control genes such as *CDKN2A* ^24^. This has prompted the development of EZH2 inhibitors that are either recently approved or are in clinical trials ^25, 26^.

Regulation of gene expression represents only one facet of EZH2 activity. Repetitive sequences that make up the majority of mammalian genomes harbour H3K27me3 and other repressive epigenetic modifications ^27–34^. These modifications repress transcription of repetitive elements and limit mobility in the host genome. Loss of epigenetic modifications such as H3K27me3 results in their upregulation. While a subset of repetitive elements has been exapted to serve the host cell, their derepression and subsequent upregulation have been linked to tumorigenesis ^35–43^. Overall, the impact of EZH2 in mediating repression of repetitive elements in normal mammalian physiology is relatively unexplored.

Pharmacologically induced upregulation of repetitive elements by inhibiting repressive epigenetic writers has been shown to elicit anti-tumor responses ^44–47^. Tumor cells treated with small molecule inhibitors against DNA methyltransferases (DNMTs) derepress the transcription of repetitive elements ^44–47^. These transcripts form secondary structures that are detected by nucleic-acid sensing pattern recognition receptors (PRRs) such as RIG-I, MDA5, and cGAS. In general, PRRs are a part of the innate immune surveillance that detect molecular patterns associated with infectious agents such as bacteria and viruses, and signals downstream to either neutralize the threat or further activate the adaptive immune system ^48, 49^. This phenomenon has been described as ‘viral mimicry’, as upregulation of repetitive elements and activation of PRRs mimics a viral infection. While viral mimicry has been demonstrated in cancer cells as a therapeutic paradigm, it is unclear if untransformed cells with normal establishment of both DNA and histone methylation to silence repeat expression are capable of viral mimicry.

Interestingly, mice with defective EZH2 recruitment to repetitive elements caused by a mutation in the retinoblastoma tumour suppressor protein (pRB) ectopically express repeats and succumb to lymphomas that often arise in the spleen and lymph nodes ^32^. To investigate the significance of EZH2 regulation of genomic repeats, we utilized pharmacological inhibition of EZH2 to investigate its acute effects on repeat regulation. Short term EZH2 inhibition with three different inhibitors induced expression of repetitive elements specifically in B cells and was accompanied by inflammation and cell death. To investigate if this effect is dependent on repeat expression, we generated triple mutant *Rigi*, *Ifih1* (MDA5), and *Cgas* mutant mice (referred to as RIC mutant) to block detection by pattern recognition receptors. In both WT and RIC mutant B cells, EZH2 inhibition induced loss of H3K27me3 at repetitive elements and increased repeat expression. Unlike WT, the RIC mutant mice failed to upregulate pro-inflammatory chemokine genes in their B cells and preserved cell viability. This gene expression program resembles EBV infected B cells suggesting it is a form of viral mimicry.

## Results

### Pharmacological EZH2 inhibition causes splenic B cell apoptosis

Constitutive loss of EZH2-mediated repression of repetitive elements in immune cells caused by mutations in *Rb1* leads to their sporadic expression and the eventual formation of lymphomas^32^. To determine the effect of acute inhibition of EZH2 in resting splenocytes, 6-8 week old mice were injected with vehicle or three different EZH2 inhibitors (GSK343, UNC1999, and EPZ6438) and spleens were harvested for histology and flow cytometry two days later (Fig. 1A). In contrast to vehicle treatment, tingible body macrophages were evident within follicles upon treatment with all three EZH2 inhibitors, suggesting B cell death and engulfment (Fig. 1B). Distinct staining for cleaved caspase 3 in spleen follicles (Fig. 1B), and the increased percentage of positively stained cells in the spleen (Fig. 1C) further suggests B cell death in response to EZH2 inhibition. Flow cytometry of splenocytes revealed a quantitative loss of CD-19+ B cells in GSK343 treated spleens and an increase in CD11b^+^Ly6C^int^Ly6G^high^ neutrophils at the same time point (Fig. 1D). Following five days of GSK343 treatment we performed H&E staining as well as IHC staining for CD68, a marker of monocytes and macrophages.

**Fig. 1.**
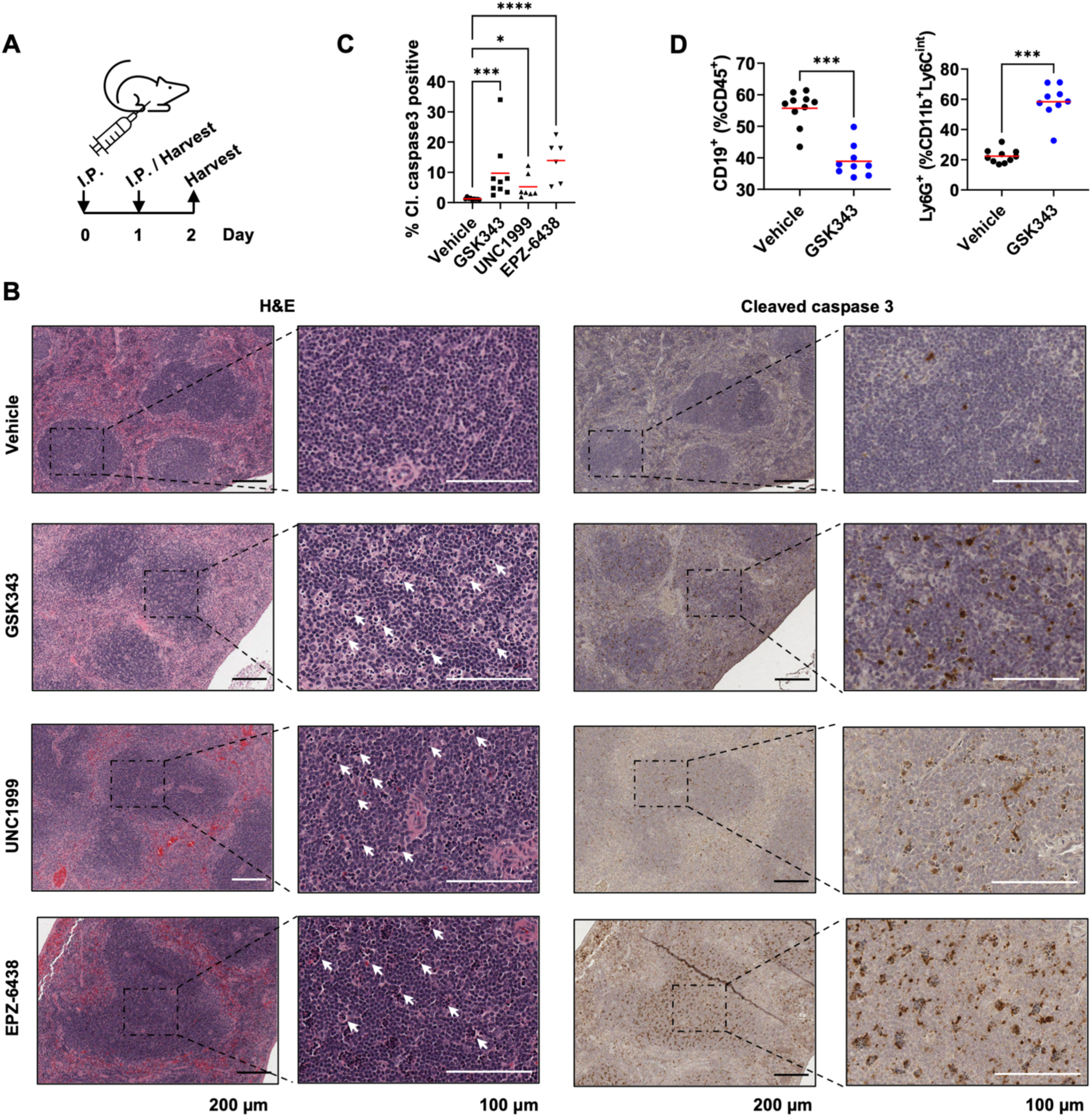
EZH2 inhibition induces B cell apoptosis and inflammation. (A) Schematic illustrating I.P. drug injection schedule for one or two-day treatments. (B) H&E staining of spleens following two days of vehicle, 100 mg/kg GSK343, 100 mg/kg UNC1999, or one day of 100 mg/kg EPZ6438. White arrows indicate tingible body macrophages (left). DAB staining for cleaved caspase 3 on consecutive spleen sections corresponding to H&E staining. (C) Percentage of cells positive for cleaved caspase 3 in the spleens from vehicle or EZH2 inhibitor treated mice. Horizontal line indicates the mean. * p < 0.05, *** p < 0.005, **** p < 0.0005 by one-way ANOVA with Krustal-Wallis multiple test correction. (D) Percentage of CD19^+^ or CD11b^+^ Ly6C^int^ Ly6G^high^ cells in the spleen following two days of vehicle or 100 mg/kg GSK343 based on flow cytometry. *** p < 0.001 by Mann-Whitney test.

This revealed that GSK343 treatment extensively reduced follicular regions of spleens and the smaller follicles that remained included notable acellular regions (Fig. S1A). CD68 staining in vehicle treated controls was largely restricted to red pulp while GSK343 induced infiltration of these cells into the follicles (Fig. S1B). Overall, these results show that targeted, short-term inhibition of EZH2 extensively disrupts splenic follicles. EZH2 inhibition causes apoptotic B cell death and the coincident arrival of neutrophils and macrophages indicates it is associated with inflammation. This was observed with three chemically distinct inhibitors, emphasizing that acute EZH2 inhibition underlies these observations.

### Unique B cell heterochromatin structure allows EZH2 inhibition to increase repeat expression

Given EZH2’s role in B lineage development ^16^, we sought to understand how its inhibition affects transcript levels in resting B cells and compare it with other resident cells of the spleen. Since EZH2 inhibition causes apoptosis *in vivo* (Fig. 1B), we treated splenic B cells *in vitro* with GSK343 and harvested before loss of viability was observed to determine the most direct impact of these inhibitors.

We separated erythrocyte-lysed splenocytes into a CD43^−^ mature splenic B cell fraction and a CD43^+^ non B-cell fraction composed of Neutrophils, T cells, and others using paramagnetic beads (Fig. 2A). We treated B cell cultures with DMSO or 1 µM GSK343 for 48 h in biological triplicates. RNA was extracted and sequenced followed by analysis with two established analysis pipelines ^50–52^ to quantify expression of repetitive elements annotated in RepeatMasker ^53^. This experiment revealed that B cells upregulated numerous repetitive elements including LINE/SINEs, LTR containing ERVs, satellite sequences, and DNA transposons (Fig. 2B, S2A). To confirm these findings with multiple EZH2 inhibitors and compare with other cells in the spleen, we carried out qRT-PCR for three classes of repeats identified as increased by RNA-seq. Figure 2C shows significant upregulation of three different classes of repetitive elements upon EZH2 inhibitor treatment of B cells (top row), but these elements were not altered by the same treatment of CD43+ splenocytes (bottom row). This data indicates repeat misexpression is a unique consequence of EZH2 inhibition in B cells and not other resident splenocytes.

**Fig. 2.**
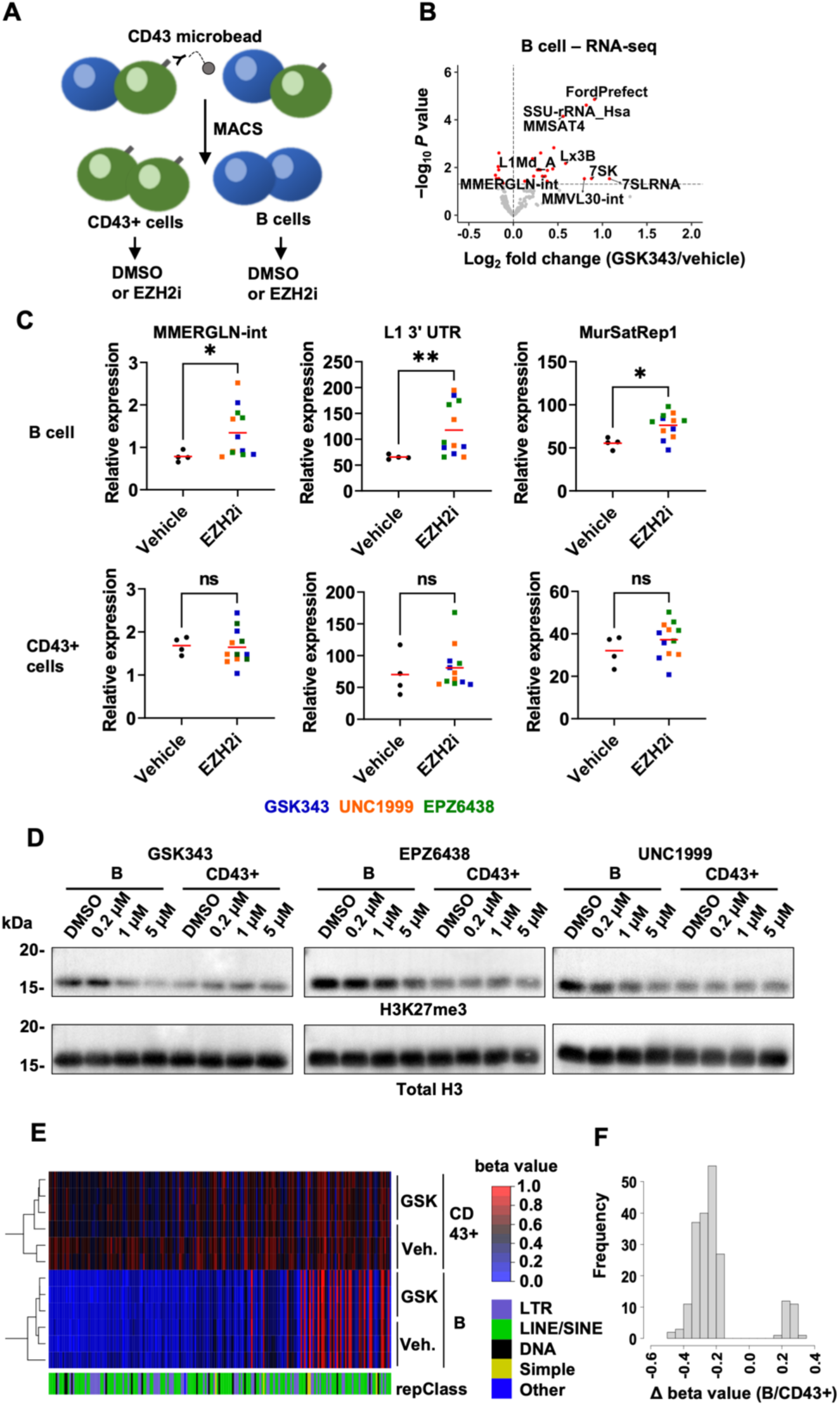
Splenic B cells are uniquely sensitive to EZH2 inhibitors. (A) Schematic describing magnetic-assisted cell sorting (MACS) to separate B and CD43+ cells from the spleen. (B) Volcano plot depicting up-or downregulated repetitive elements in purified splenic B cells treated with DMSO or 1 µM GSK343 for 48 h in culture. (C) qRT-PCR of indicated repetitive elements in splenic B cells (top row) or non-B cells (bottom row) treated with DMSO, GSK343, UNC1999 or EPZ6438. * p < 0.05, ** p < 0.005, **** p <0.0001 by Mann-Whitney test. (D) Western blots of H3K27me3 and total H3 from histone extracts of B or CD43+ cells treated with DMSO or increasing concentrations of indicated EZH2 inhibitors for 48 h in culture. (E) Heatmap of DNA methylation probe beta values at top 200 differentially methylated probes annotated with repetitive elements by repClass between DMSO-treated B and CD43+ cells. (F) Histogram of difference in beta values (DMSO-treated B vs. CD43+ cells) among top 200 differentially methylated probes shown in E.

To understand the regulation of B cell heterochromatin better, cultures of B and CD43+ cells were incubated for 48 hr with increasing concentrations of EZH2 inhibitors (Fig. 2D). Histones were extracted and analyzed by western blotting for H3K27me3 to directly investigate the effect of EZH2 inhibition. This demonstrated that B cells have higher baseline H3K27me3 compared to CD43+ cells (Fig. 2D). In addition, loss of H3K27me3 was observed with increasing inhibitor concentration in B cells with all three EZH2 inhibitors (Fig. 2D). Cells in the CD43+ fraction did not show an obvious reduction in H3K27me3 levels in response to any of these treatment conditions. This indicates a unique reliance on EZH2 for the maintenance of heterochromatin in B cells that is distinct from other cell types in the spleen.

Repression of repeat expression is also known to be mediated by DNA methylation, therefore we also investigated its status in B and CD43+ cells. We performed a genome-wide DNA methylation microarray on DNA extracted from vehicle or GSK343 treated B and CD43+ cells. Differential beta value analysis between vehicle treated B vs. CD43+ cells revealed that probes annotated as representing repetitive elements had low levels of DNA methylation in B cells compared to CD43+ splenocytes (Fig. 2E). As expected, GSK343 treatment did not affect DNA methylation in either cell fraction. Only a small proportion of differentially methylated repeat probes were more methylated in B cells compared to the NB fraction (Fig. 2F). By comparison, DNA methylation levels observed at probes annotated for promoters, genes or CpG islands displayed much greater similarity between B and CD43+ populations (Fig. S2B). These data demonstrate low level DNA methylation at repeat elements uniquely in B cells.

Taken together, these experiments show that splenic B cells specifically upregulate transcription of repetitive elements upon EZH2 inhibition. This corresponds with loss of H3K27me3 and constitutive low levels of DNA methylation at repetitive elements. This unique B cell heterochromatin helps explain why systemic EZH2 inhibitor treatment causes a focused disruption of splenic follicles and B cell death. An intriguing interpretation of these experiments is that EZH2 inhibition may stimulate an inflammatory response through misexpressed repeats in B cells that leads to their elimination.

### Pattern recognition receptors are required for EZH2 inhibitor-induced B cell death in the spleen

Ectopic expression of repetitive elements and subsequent inflammatory signaling are described as a state of viral mimicry^54, 55^. Transcripts from repetitive elements form secondary structures that mimic those of viral replication and transcription. Cytosolic nucleic acid sensing pattern recognition receptors (PRRs) such as RIG-I, MDA5 and cGAS bind to dsRNA/DNA to activate a signaling cascade that upregulates inflammatory gene expression. Notably, others have shown that these cytosolic PRRs are mechanistically required for DNMT or EZH2 inhibition-induced anti-tumour immune responses. We sought to determine if activation of pattern recognition receptors in splenic B cells underpins the inflammatory response and cell death following EZH2 inhibition.

We created a loss of function mouse model in which PRR genes are disrupted. Briefly, three gRNAs, each targeting a coding exon for one of *Rigi*, *Ifih1* (MDA5), and *Cgas* were microinjected with a Cas9 expressing mRNA into single-cell zygotes to simultaneously mutate all three genes and disrupt their coding capacity ^56^ (Fig. 3A). The resulting founder pups were bred together to create biallelic F1 mice that were characterized for their mutant alleles (Fig. S3A). Mutants were identified using a restriction fragment length polymorphism assay to identify indels in the three genes in F1 mice (Fig. S3B,C). All mutant alleles in our triple mutant colony were sequenced to characterize how they disrupt their respective gene (Table S1). Lastly, we sought to identify any off-target mutations created by the gRNAs in the F1 mice that we bred for experiments. We sequenced the two highest ranked coding and non-coding, *in silico* predicted ^57^, off-target loci per gRNA (six in total) in the three F1 mice. Out of 36 possible alleles, we only found two, one-bp deletions in non-coding regions (Table S2) confirming high fidelity of targeting by this strategy. All triple mutant *Rigi*, *Ifih1*, and *Cgas* mice (henceforth called RIC mutant) used in this study are descendants from these three F1 animals and were compared against C57BL/6NCrl controls.

**Fig. 3.**
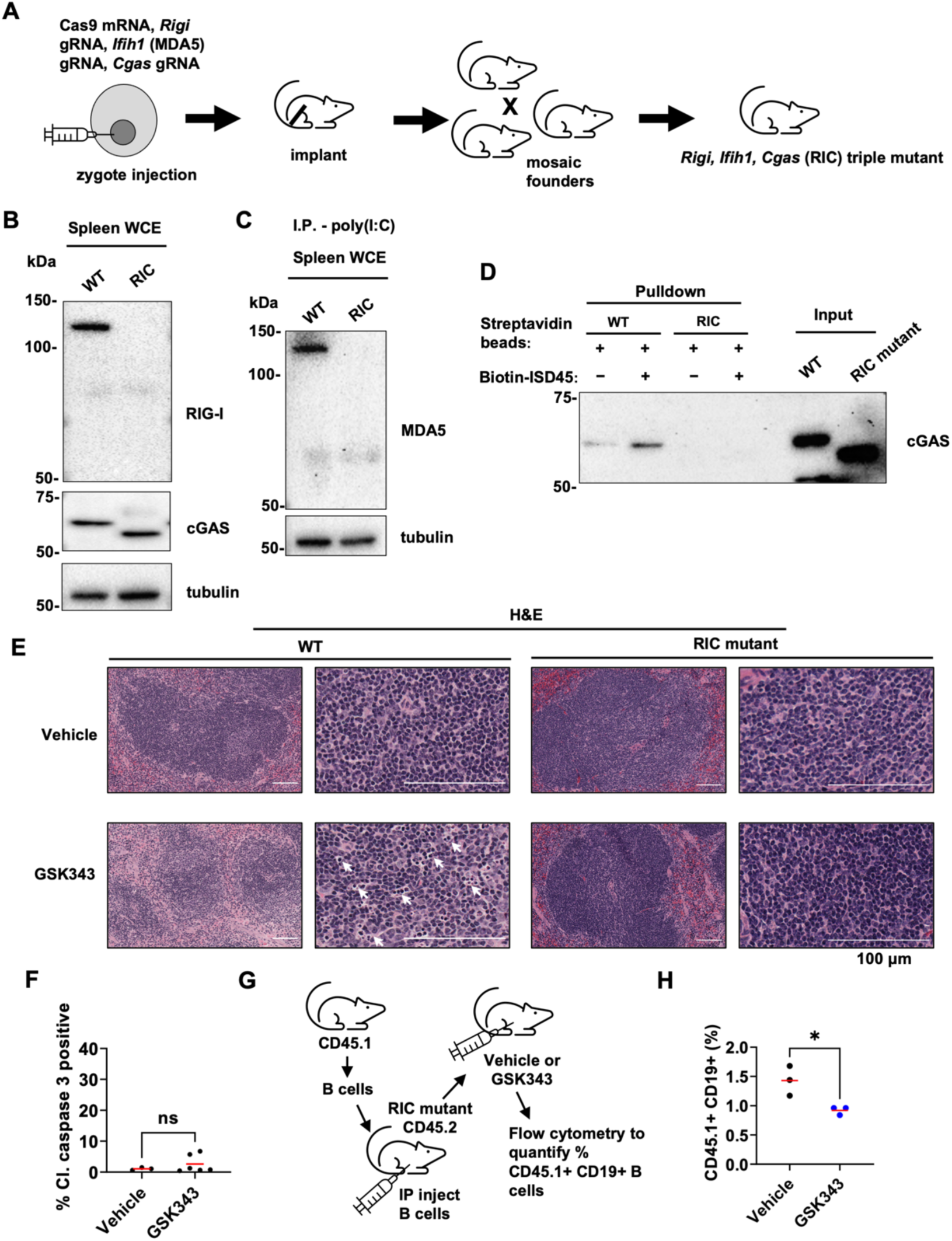
*Rigi/Ifih1/Cgas* (RIC) triple mutant mice are resistant to EZH2 inhibitor killing of B cells. (A) Schematic highlighting the key steps of generating RIC triple mutant mice. gRNAs and Cas9 mRNA are microinjected into zygotes and transplanted into a surrogate female. Resulting mosaic founders are bred together to generate bona fide triple mutant mice. (B) Western blots of RIG-I and cGAS from whole cell extracts of WT and RIC mutant splenocytes. (C) Western blots of MDA5 from whole cell extracts of WT and RIC mutant splenocytes harvested from mice injected with poly(I:C). (D) Western blots of cGAS pulldown with biotinylated ISD45 probe and streptavidin beads from whole cell extracts of WT and RIC mutant splenocytes. (E) H&E staining of the spleens following 2 days of vehicle or 100 mg/kg GSK343 in WT or RIC mutant mice. White arrows indicate tingible body macrophages. (F) Percentage of cells positive for cleaved caspase 3 in the spleens from vehicle or GSK343 I.P. injected RIC mutant mice. Horizontal line indicates the mean. ns p > 0.05 by Mann-Whitney test. (G) Schematic describing adoptive transfer of CD45.1 WT B cells into CD45.2 RIC mutant mice, followed by vehicle or GSK343 injections. (H) Percentage of CD45.1+ CD19+ adoptively transferred, WT B cells in RIC mutant spleens upon vehicle or GSK343 injections. * p < 0.05 by unpaired Student’s t-test.

We characterized expression from these mutant alleles by western blotting. RIC mutant splenocytes completely lost RIG-I and MDA5 expression (Fig. 3B,C), confirming that *Rigi* and *Ifih1* mutations are null alleles. The truncated cGAS detected by western blotting (Fig. 3B) agrees with the 48-bp deletions found in two distinct *Cgas* alleles (Fig. S3D,E). These N-terminal, in frame deletions, encode functionally inactive cGAS as it fails to bind a known dsDNA target, ISD45, in a streptavidin-biotin pulldown assay^58^ (Fig. 3D). Furthermore, streptavidin alone has been shown to bind and activate cGAS, and truncated cGAS has also lost this interaction ^59^ (Fig. 3D).

To understand the role of PRRs in the response to EZH2 inhibition, systemic treatment of RIC mutant mice with GSK343 was performed and spleens were examined after two days. Infiltration of tingible body macrophages was less prominent in RIC mutant follicles (Fig. 3E). Furthermore, the percentage of cells positive for cleaved caspase 3 was not significantly increased upon GSK343 treatment in RIC mutants (Fig. 3F). These observations suggest that the functional loss of the cytosolic PRRs in the RIC mutant largely abrogates the cell death phenotype in spleens of GSK343 treated mice. However, all cells in the RIC mutant mouse are disrupted for PRRs and not just their B cells. PRR mutations may affect other cell functions, possibly those involved in an inflammatory response that could prevent B cell death in response to EZH2 inhibitors. To rule out this possibility we isolated B cells from C57BL/6 CD45.1 donors and transferred them to CD45.2 RIC mutant recipients where they were treated with GSK343 for 2 days (Fig. 3G). Flow cytometry to identify CD45.1 B cells from the spleens of these mice revealed that GSK343 treatment diminishes WT B cells in a RIC mutant host. This data argues that the mutant host has no ability to block B cell death induced by inflammation from EZH2 inhibitors.

Using CRISPR-Cas9, we generated triple mutant RIC mice that are deficient for PRR function. The functional loss of the cytosolic PRRs blocks EZH2 inhibitor induction of B cell death in the spleen. Adoptive transfer experiments demonstrate that WT B cells are lost following GSK343 treatment of RIC mutant hosts. Collectively, this suggests that PRRs respond to misexpressed repeats in B cells following EZH2 inhibitor treatment, causing inflammation and cell death.

### EZH2 inhibition induces H3K27me3 loss at repetitive elements in B cells

We investigated the effect of EZH2 inhibition at the chromatin level in isolated WT and RIC mutant B cells. We performed ChIP-seq for H3K27me3 in DMSO or GSK343 treated cells in biological duplicates. We obtained high quality sequencing libraries from input controls and H3K27me3 associated fragments (Fig. S4A,B), we then identified H3K27me3 peaks using MACS2 and quantified read fragment enrichment at those peaks. As expected, most high-scoring peaks found in vehicle treated samples were shared by WT and RIC mutants (Fig S4C), confirming that the baseline locations of H3K27me3 deposition is independent of functional expression of PRRs. We then compared loss of H3K27me3 upon GSK343 treatment with their respective vehicle treated genotype controls. Fig. 4A depicts normalized read fragment enrichment where each row represents a scaled peak length with 1 kb flanking each end. The sum of the rows represents all the peaks identified in vehicle-treated, control ChIP samples for each genotype (WT and RIC mutant). At these baseline H3K27me3 peaks, GSK343 decreased enrichment in both WT and RIC mutants (5^th^ and 6^th^ vs. 7^th^ and 8^th^ columns). Next, we sought to determine which genomic features were associated with these H3K27me3 peaks. We annotated the peaks found in each ChIP sample (pooling biological replicates together) based on their proximity to known genes. The absolute fold decrease in peak count was the greatest in intronic and intergenic regions for both WT and RIC mutant upon GSK343 treatment (Fig. 4B,C). Intronic and intergenic regions contain repetitive elements and the absolute number of repetitive elements that intersect with peaks was similarly decreased upon GSK343 treatment in both genotypes (Fig. 4D,E, S4D). In addition, we confirmed that H3K27me3 enrichment at peaks intersected with specific repeat element families identified by RNA-seq and that these were decreased upon GSK343 treatment in both genotypes (Fig. 4F).

**Fig. 4.**
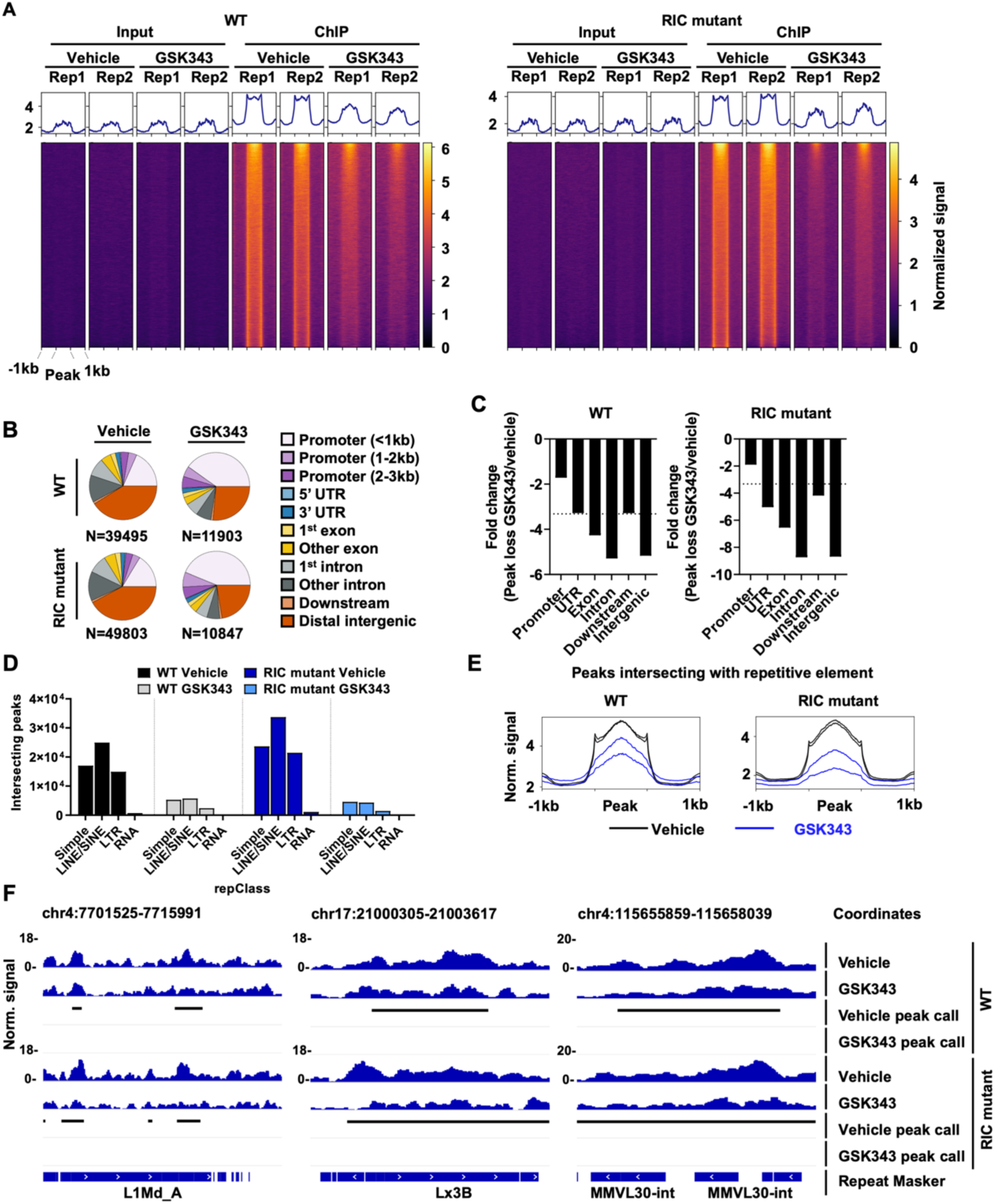
GSK343 induces loss of H3K27me3 at repetitive elements in splenic B cells. (A) Heatmap of input and H3K27me3 ChIP-seq read enrichment at all peaks called in DMSO treated samples. Splenic B cells were treated with either DMSO or 1 µM GSK343 for 48h. Enrichment was quantified as reads per genomic content (RPGC). Each row represents a scaled DMSO peak location with 1 kb flanking each end. Rows are sorted by decreasing enrichment. (B) Distribution of peaks amongst the indicated genomic features in each sample. (C) Fold-change of the number of called peaks annotated with indicated genomic features. Horizontal dotted lines indicate the average overall fold change. (D) Number of repetitive elements in each indicated repClass that intersect with called peaks in each sample condition shown. (E) Enrichment profiles of H3K27me3 ChIP-seq reads at peaks called in DMSO or GSK343 treated samples intersecting with repetitive elements. Each box shows the average profile of scaled repetitive elements with 1 kb flanking each end. Biological duplicates for each treatment condition are shown as separate curves. (F) Three genome track views showing normalized coverage (RPGC) of H3K27me3 signal track for indicated sample conditions at repetitive elements. Horizontal bars indicate either peak calls or repetitive element annotations.

These ChIP experiments show that EZH2 inhibition decreases H3K27me3 preferentially at repetitive elements. This is observed in both WT and RIC mutant B cells. It confirms that inactivation of cytosolic PRRs has no bearing on H3K27me3 containing heterochromatin or the effect of EZH2 inhibition on H3K27me3 reduction. These findings suggest PRRs mediate effects downstream of H3K27me3 loss following EZH2 inhibition in B cells.

### Cytosolic PRRs are required for pro-inflammatory gene expression upon EZH2 inhibition

Since GSK343 induces comparable loss of H3K27me3 in both WT and RIC mutant B cells, we next investigated its effects on the transcriptome. WT and RIC mutant B cells were treated with DMSO or GSK343 as before, RNA was extracted, and we performed RNA-seq to identify differentially expressed repetitive elements, genes, and significantly enriched gene sets.

This revealed that the most differentially upregulated genes upon EZH2 inhibition were correlated with loss of H3K27me3 at their promoters (Fig. 5A), and this effect was observed in both genotypes. Consistent with similar effects by GSK343 on H3K27me3 between WT and RIC mutant mice, upregulation of many of the same repeats also took place in RIC mutants (Fig. 5B, S5A,B). From these analyses we conclude that EZH2 inhibition leading to H3K27me3 loss causes similar increases in gene and repeat transcript levels in WT and RIC mutants.

**Fig. 5.**
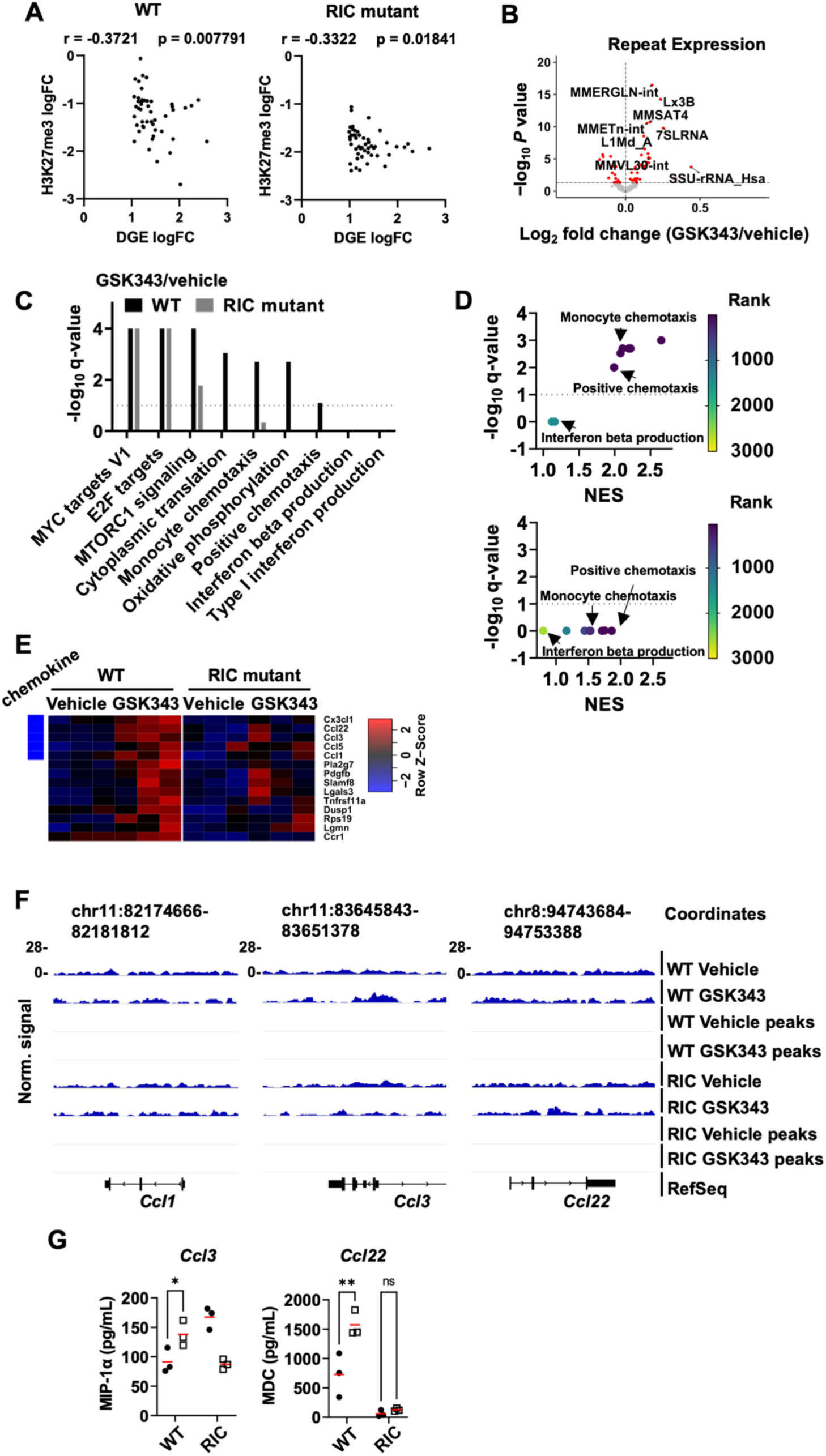
Cytosolic PRRs are required for GSK343-induced inflammatory signaling in B cells. (A) Scatter plot showing a negative correlation between loss of H3K27me3 near promoters and upregulation of nearby genes in WT and RIC mutant splenic B cells. Non-parametric Spearman’s correlation. (B) Volcano plot depicting up-or downregulated repetitive elements in RIC mutant splenic B cells treated with DMSO or 1 µM GSK343 for 48 h in culture. (C) Bar plot depicting adjusted P values (FWER) of GSEA of Gene Ontology (GO) biological processes and Hallmark gene sets. Horizontal dotted line indicates P value cut-off at 0.1. (D) Net enrichment scores (NES), q-value (FWER) and rank among gene sets (colour legend) for chemotaxis or IFN-related gene sets for WT (top) and RIC mutant (bottom). Horizontal dotted line indicates P value cut-off at 0.1. (E) Expression heatmap of genes annotated in “monocyte chemotaxis” gene set in WT and RIC mutant splenic B cells. Expression is shown as a Z-score of the mean of each row and chemokine genes are indicated. (F) Genome track view showing normalized coverage (RPGC) of H3K27me3 signal tracks at genes encoding chemokines in “monocyte chemotaxis” GO pathway. (G) Quantification of MIP-1α (*Ccl3*), MDC (*Ccl22*) and IFN-β from cell culture supernatant of WT and RIC mutant splenic B cells treated with DMSO, 0.1 or 1 µM GSK343 for 48 h in culture. * p < 0.05, *** p < 0.001 by two-way ANOVA with Dunnett’s multiple test correction.

To understand the consequences of PRR loss in response to EZH2 inhibition, we searched for differences in gene expression that were not the consequence of direct regulation by H3K27me3. Most genes that were significantly up or downregulated in GSK343 treated WT cells showed a similar change in RIC mutants (Fig. S5C), and many gene sets were commonly upregulated in both WT and RIC mutants (Fig. 5C, S5D, Table S3). However, RIC mutants showed a smaller absolute number of genes with altered expression compared to WT (Fig. S5C). Furthermore, several pathways that were significantly enriched in WT, were missing in RIC mutants (Fig. 5C). In particular, pathways related to chemotaxis were enriched in GSK343 treated WT samples but not RIC mutants, suggesting a potential source of inflammatory signaling dependent on PRRs. We note that interferon signaling, described in previous studies of viral mimicry upon epigenetic inhibition ^44–47^, is not activated by GSK343 in B cells (Fig. 5C,D). Upregulation of monocyte chemotaxis genes were more consistent and robust in WT compared to RIC mutants (Fig. 5E). Notably, several of these genes were secreted chemokines with pro-inflammatory functions. Importantly, upregulation of these chemokines in WT, but not RIC mutant B cells, was not associated with direct loss of H3K27me3 as evidenced by the absence of peaks at these genes under all experimental conditions (Fig. 5F). This suggests that their upregulation is mediated by GSK343-induced repetitive element expression, activation of cytosolic PRRs and subsequent signaling, and not direct derepression of gene expression by EZH2 inhibition.

Consistent with gene expression data, detection of secreted cytokines by array analysis demonstrates RIC dependent upregulation of MIP-1α (encoded by *Ccl3*) and MDC (encoded by *Ccl22*), but not IFNβ upon GSK343 treatment (Fig. 5G). This is consistent with our finding that chemokines, but not interferons, are activated by GSK343 treatment. In sum, these data indicate that the cytosolic PRRs are required to upregulate pro-inflammatory cytokine signalling and chemokine production upon EZH2 inhibition with GSK343 and this effect is blocked in RIC mutants. This observation is not attributable to lack of EZH2 inhibition in RIC mutants because upregulation of repetitive elements upon GSK343 treatment is retained. We suggest that in WT B cells, EZH2 inhibition activates PRRs that stimulates expression and secretion of chemokines to induce inflammation and cell death.

### EZH2 inhibition resembles B cell infection by Epstein-Barr Virus

To understand if EZH2 inhibition induced inflammation and B cell death represent synthetic chemical effects or are triggering a natural physiological response, we compared our findings with B cells following viral infection. We performed the same GSEA on existing RNA-seq data of Epstein-Barr Virus (EBV) infected human resting B cells ^60^. This revealed that monocyte chemotaxis was a commonly shared and highly ranked gene expression category from both EBV infection and EZH2 inhibition (Fig. 6A,B). Furthermore, we found that the monocyte chemotaxis gene set was significantly enriched in EBV infected cells compared to control at five out of six time points following infection (Fig. 6C). In addition, upregulation of *Ccl22, Ccl3,* and *Ccl5* were comparable between GSK343 treated WT B cells and EBV infected human B cells, while GSK343 treated RIC mutant B cells did not significantly upregulate them (Fig. 6D). These similarities in gene expression of pro-inflammatory chemokines suggest that EZH2 inhibition induces repeat expression in B cells that mimics the response to an EBV infection. Based on this similarity we describe EZH2 inhibition in B cells as a viral mimicry response and emphasize that its mechanism of action is clearly distinct from previously reported viral mimicry that is highly dependent on interferon signaling.

**Fig. 6.**
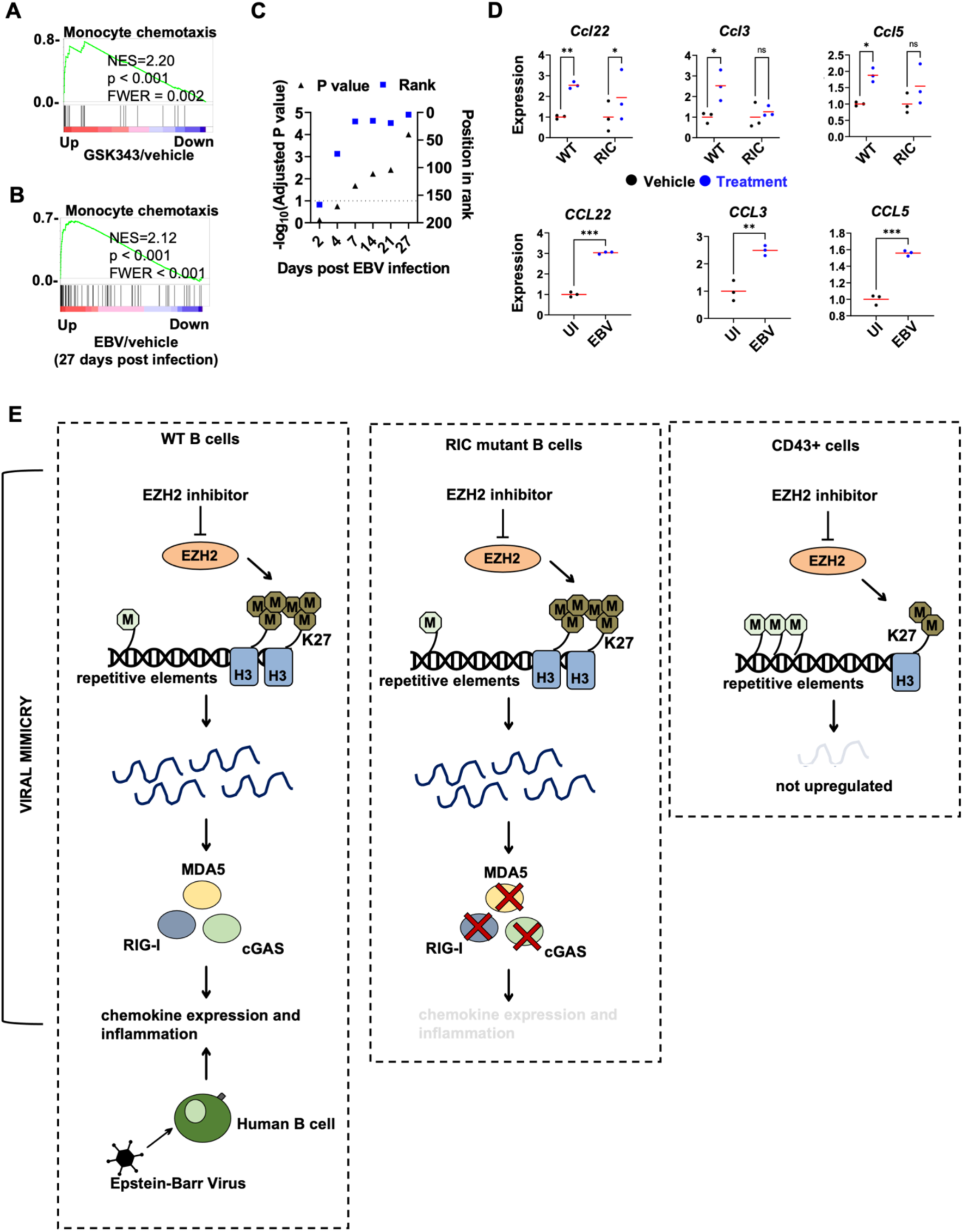
EZH2 inhibition mimics B cell infection by Epstein-Barr Virus. (A) GSEA plot of “monocyte chemotaxis” GO gene set significantly enriched in GSK343-treated splenic B cells compared to vehicle. (B) GSEA plot of “monocyte chemotaxis” GO gene set significantly enriched in EBV-infected human resting B cells compared to control. (C) Adjusted P values (FWER) and position in ranked list (out of all tested gene sets) of “monocyte chemotaxis” GO biological process pathway at indicated days post EBV infection in human resting B cells. Horizontal dotted line indicates P value cut-off at 0.1. (D) Expression of chemokines that are significantly upregulated in GSK343 treated WT B cells (compared to vehicle) and EBV infected human resting B cells (compared to control). Horizontal red bars represent the mean. Top row: * p < 0.05, ** p < 0.005 by two-way ANOVA with Sidak’s multiple test correction. Bottom row: ** p < 0.005, *** p < 0.001 by unpaired Student’s t-test. (E) A proposed model of viral mimicry upon EZH2 inhibition in splenic B cells.

## Discussion

Our work demonstrates a central role for PRRs in responding to EZH2 inhibition in mammals. We revealed that EZH2 inhibition uniquely activates repetitive element expression in B cells. Repeat misexpression is detected in WT and RIC mutant B cells treated with EZH2 inhibitors, indicating that the functional loss of the PRRs does not compromise the initial, on-target effects of EZH2 inhibition. In contrast, loss of PRR function in RIC mutant B cells failed to activate expression of pro-inflammatory cytokines and inflammation and cell death were blocked. Based on these findings, we propose the signaling model in Fig. 6D that highlights our discovery of a key aspect of EZH2 chemical inhibition that is a viral mimicry response leading to inflammation and B cell death. This has important implications for H3K27Me3 in heterochromatin formation and EZH2 inhibitor use and it suggests new applications for this class of therapeutic.

Our H3K27me3 ChIP-seq data from B cells reveals a striking preference for EZH2 inhibition causing decreased H3K27me3 in intronic and intergenic regions in comparison with proximal promoters. Furthermore, our work reveals that there is extensive overlap of H3K27me3 intersecting with repetitive elements and EZH2 inhibitors induces their expression. In addition, we demonstrate that B cell repeat sequences lack CpG methylation. This unique combination of heterochromatin characteristics largely explains why B cells are susceptible to viral mimicry in response to EZH2 inhibitors. We are unaware of other cell types that possess this combination of chromatin characteristics as we did not detect this striking cell death and inflammation phenotype in other major organs or tissues.

Our data highlights the similarity between Epstein-Barr Virus infection and the response to EZH2 inhibitors that is best appreciated by the similarity of chemokines whose expression they both activate. Expression of *Csf1*, *Ccl22* (MDC), *Ccl1*, *Ccl3*, and *Ccl5* are all induced upon infection of B cells ^61, 62^, and these chemokines are all activated in a PRR dependent manner by EZH2 inhibitors in our experiments. The mechanism by which B cell death is induced likely depends on these chemokines, although the mechanism is also unclear in the response to EBV infection. Some studies indicate that EBV infected cells are targeted by T and NK cells following infection ^63, 64^, and this is consistent with the inflammation and apoptotic death we observe following EZH2 inhibitor treatment. It is also possible that a more direct mechanism of apoptotic induction is responsible, although transcriptional changes found in WT B cells and missing in RIC B cells offers no obvious mechanism. Regardless of the precise explanation for B cell death, this viral mimicry that resembles EBV infection reveals a distinct pathway not observed in prior studies of interferon activating paradigms described in cancer cells.

The rationale for generating EZH2 inhibitors is to counteract overactive H3K27me3 deposition that arises from overexpressed or mutationally activated EZH2. Our study reveals the consequence of inhibition of endogenous, wild type EZH2. We suggest important considerations and potential applications for EZH2 inhibitors in the future. First, short term treatment with these inhibitors is capable of inducing inflammation and depleting B cells. Recent studies using other EZH2 inhibitors, DZNep or GSK126, to treat mouse models of systemic lupus erythematosus demonstrated phenotypic improvement in a number of measures of disease pathology ^65, 66^. These studies rationalized use of EZH2 inhibitors based on high level EZH2 expression even though this isn’t observed in all patients or relevant immune cell types. Our work demonstrates activation of viral mimicry in B cells after only brief treatment and suggests an alternate explanation to reduced B cells in these models. Secondly, the specificity of viral mimicry related immune stimulation in cancer treatment is based on the concept that cancer cells have precarious silencing of repeats. Epigenetic alterations inherent to cell transformation alter heterochromatin such that inhibitors of DNA methyltransferases selectively cause repeat expression in cancer cells but not normal somatic cells. Our work suggests that EZH2 inhibitors may be deployed based on a similar logic whereby low levels of DNA methylation are acquired through transformation and cells are susceptible to EZH2 inhibition even without overexpression of EZH2 ^45^. Our findings also raise the question of whether the activation of inflammatory signaling upon EZH2 inhibition in B cells from prolonged treatment with this class of inhibitor can lead to chronic inflammation. Overall, our work reveals a new effect of EZH2 inhibitors in immune function that is likely to have a significant impact on the use of these agents in clinical applications.

## Acknowledgements

The authors are grateful to Drs. Nathalie Berube and Lisa Cameron for advice in preparing this manuscript. S.J.K was funded by Cancer Research and Technology Transfer training program and an Ontario Graduate Scholarship. This work was supported by a grant from the Canadian Institute of Health Research (PJT-156014) to F.A.D. The authors wish to thank numerous colleagues at the London Regional Cancer Program for suggestions and encouragement in the course of this work.

## Author contributions

S.J.K and F.A.D. designed the experiments and wrote the manuscript. P.K.K. performed histological analysis in Fig. 1B-D and Fig. 2F. R.P.D provided guidance on MACS isolation of B cells and analysis of immune function. S.J.K. performed all experiments and data analysis.

## Methods

### Resource availability Lead contact

Further information and requests for resources and reagents should be directed to and will be fulfilled by the lead contact, F.A.D.

## Materials availability

RIC mutant mice generated in this study are available from the lead contact upon request.

### Data and code availability

All high-throughput sequencing data have been deposited at GEO (GSE198232) and are publicly available as of the date of publication. The accession number is listed in the key resources table. RNA-seq data of EBV infected human resting B cells was obtained from GEO under GSE125974 ^60^. Original Western blot images have been deposited at Mendeley and are publicly available as of the date of publication. This paper does not report original code. Any additional information required to reanalyze the data reported in this paper is available from the lead contact upon request.

### Experimental model and subject details

C57BL/6NCrl (Charles River, #027) and RIC triple mutant mice (described below) were housed and monitored according to institutional animal use guidelines (protocol number 2020-039). Mice were given access to standard chow and water ad libitum and exposed to 12 h light/dark cycles in a pathogen-free exclusion facility. For I.P. injections and splenic B cell isolation, littermates 6-8 week old mice of both sexes were randomly assigned to experimental groups within each genotype.

## Method details

### Intraperitoneal injections

For EZH2 inhibition *in vivo*, 6-8 week old mice were I.P injected daily with 100 mg/kg GSK343 (Tocris, #6128) for two or five days, UNC1999 (Cayman Chemical, #14621) for two days, or EPZ6438 (SelleckChem, #S7128) for one day in 20 % (w/v) Captisol (Captisol, San Diego), pH 4.5 (with 1N acetic acid). Control mice were I.P injected daily with 20 % (w/v) Captisol (Captisol, San Diego), pH 4.5 (with 1N acetic acid) for two or five days. Mice were sacrificed, and the spleen and bone marrow were harvested for analysis. For poly(I:C) (Millipore-Sigma, #P1530) treatment, 6-8 week old male or female WT and RIC mutant mice were I.P injected with either 100 µg poly(I:C) or PBS. The next day, the mice were sacrificed, and the spleen was harvested for analysis.

### Immunohistochemistry

Spleens were fixed in 10% neutral buffered formalin for 48 h prior to paraffin embedding. Routine H&E staining was performed. For immunohistochemistry, tissue sections were incubated in the following solutions for 3 min each: 100% xylene, 100% xylene, 100% ethanol, 100% ethanol, 95% ethanol, 70% ethanol. Antigen retrieval was performed in citrate (10 mM sodium citrate, pH 6.0) or Tris (10 mM Tris, 1 mM EDTA, pH 9.0) buffer for CD68 (Abcam, #125212, 1:100), and cleaved caspase 3 (CST, #9661, 1:400) staining, respectively. Slides were incubated with the primary antibodies in a humidified chamber overnight at 4 °C. Goat anti-rabbit biotinylated IgG (VectorLabs, #BA-1000-1.5), peroxidase-streptavidin (VectorLabs, #SA-5704-100) and DAB subtrate kit (VectorLabs, #SK-4100) were used the next day at RT to develop the staining.

Mayer’s hematoxylin was used to counterstain, then washed with tap water to destain. Slides were dehydrated and mounted with a coverslip and mounting medium (VectorLabs, #H-5000). DAB positive cell detection based on a maximum threshold in QuPath/0.3.2 was used to identify the percentage of cleaved caspase 3 positive cells.

### Cell culture

For splenocyte culture, spleens were harvested from untreated WT and RIC mutant mice, gently homogenized with a syringe plunger and passed through a 40 µm mesh filter (Fisher Scientific, #08-771-1). Filtered cells were centrifuged at 300 x g for 10 min at 4 °C. The cell pellet was resuspended in ACK lysis buffer (150 mM NH_4_Cl, 10 mM KHCO_3_, 0.1 mM EDTA, pH 7.2) for 4 min at RT to lyse erythrocytes. Remaining splenic lymphocytes were washed twice with FACS buffer (5% BSA, 2 mM EDTA in PBS) and incubated in RPMI-1640 (Wisent) supplemented with 10% (v/v) FBS, 55 µM 2-ME, 2 mM L-glutamine, penicillin and streptomycin at 37 °C in 5% CO_2_.

To isolate splenic B cells and non-B cells, washed splenic lymphocytes (5 x 10^7^) were resuspended in 450 µL FACS buffer and stained with 50 µL CD43 microbeads (Miltenyi Biotec, #130-049-801) on ice for 30 min. Cells were washed once, resuspended in 500 µL FACS buffer and passed through an LD column in a VarioMACS separator (Miltenyi Biotec, #130-042-901, #130-090-282) as per manufacturer’s recommendation. CD43 microbead-labelled non-B cells were eluted from the column by applying the plunger. Purified B cells or non-B cells were incubated in RPMI-1640 supplemented as above plus 2 ng/mL IL-4 and BAFF (BioLegend, #574302, #591202) at 37 °C in 5% CO_2_. These cells were treated with DMSO, GSK343 (Tocris #6128), UNC1999 (Cayman Chemical, #14621) or EPZ6438 (SelleckChem, #S7128) for 48 hr in culture. Cell viability was measured by trypan blue exclusion assay and quantified by Countess II (ThermoFisher).

### Infinium mouse methylation array

Genomic DNA was purified from B or non-B cells using Monarch genomic DNA purification kit (NEB, #T3010). Further sample processing and mouse methylation array (Illumina) were performed by The Centre for Applied Genomics, The Hospital for Sick Children, Toronto, Canada. “Combined rank” metric in RnBeads/3.17 was used to generate an ordered list of differentially methylated probes between DMSO-treated B and non-B cells^67^. In addition to default probe annotations (cpgislands, genes and promoters), a custom repClass annotation was generated from RepeatMasker (downloaded from UCSC table browser) and applied to the probes. Beta values of such annotated, differentially methylated probes were represented as heatmaps (gplots /3.1.3).

### RNA extraction, qRT-PCR and sequencing

Splenocytes or splenic B cells (5 x 10^6^ cells/mL) were treated with either DMSO or 1 µM GSK343 for 48 h. RNA was harvested using Monarch Total RNA miniprep kit (NEB, #T2010) and residual genomic DNA was digested by treating 1 µg total RNA with 1 U DNaseI (ThermoFisher, #18068015) for 15 min at RT. DNaseI was then inactivated by adding EDTA and incubating at 65 °C for 10 min. For qRT-PCR, RNA was then reverse-transcribed into cDNA with iScript Supermix (Biorad, #1708840) and diluted five-fold with H_2_O. PCR was performed with iQ SYBR Green Supermix (Biorad, #1708882) on CFX96 (Biorad). All primer sequences are described in Table S4 ^68–70^. For sequencing, DNaseI-treated RNA was purified with Monarch RNA cleanup kit (NEB, #T2040). rRNA depletion and library preparation were performed with VAHTS total RNA-seq library prep kit (GeneBio, #NR603-01). Libraries were pooled and sequenced on NextSeq 500 at the London Regional Genomics Center with a high output 75 cycle kit to yield single-end 75-bp reads.

### RNA sequencing analysis

Demultiplexed FASTQ files were downloaded from BaseSpace. RepEnrich2 and featureCounts were used to quantify repetitive element expression ^50–52^. Briefly, bowtie2/2.4.2 ^71^ was used to map reads to mm10 with default settings. Resulting sam files were converted to bam files with samtools/1.12 ^72^. RepeatMasker track for mm10 was filtered to remove simple repeats, then used to build a pseudogenome with RepEnrich2 subcommands. Fractional count tables were imported to RStudio running r/3.6.3. Alternatively, STAR/2.5.2b ^73^ and featureCounts were used with recommended settings ^52^ with the filtered RepeatMasker track to generate count tables. For gene expression quantification, reads were mapped to mm10 genome with STAR/2.7.8a and resulting sam files were converted to sorted, indexed bam files with samtools/1.12. HTSeq/0.11.0 ^74^ was used to assign mapped reads to GENCODE mouse M22 comprehensive gene annotation ^75^.

To identify differentially expressed repetitive elements or genes, edgeR/3.28.1 ^76^ was used. Volcano plots, heatmaps and Venn diagrams were generated with EnhancedVolcano, heatmap.2 and VennDiagram ^77^. GSEA was performed as recommended ^78^.

### ChIP sequencing

WT and RIC mutant splenic B cells (5 x 10^6^ cells/mL, 7.5 x 10^6^ cells total) were treated with either DMSO or 1 µM GSK343 for 48 h. Cells were washed with PBS and resuspended in 1 mL 1% (v/v) formaldehyde in PBS to fix chromatin-protein complexes. After 5 min, the reaction was quenched by adding glycine to a final concentration of 0.125 M. Fixed cells were washed twice with PBS and incubated on ice for 10 min in lysis buffer 1 (10 mM HEPES pH 6.5, 10 mM EDTA, 0.5 mM EGTA, 0.25% Triton X-100). The suspension was centrifuged at 600 x g for 5 min at 4 °C to isolate the nuclei. They were washed twice in wash buffer (10 mM HEPES pH 6.5, 1 mM EDTA, 0.5 mM EGTA, 200 mM NaCl) and resuspended in lysis buffer 2 (50 mM Tris pH 8.0, 1 mM EDTA, 0.5% Triton X-100, 1% SDS) to 7.5 x 10^6^ cell/200 µL in sonication tubes. Chromatin was sonicated to 100-600 bp fragments by four cycles of 30 s ON and OFF in Bioruptor Pico (Diagenode, #B01060010). Debris was cleared by centrifugation at 16 000 x g for 30 min at 4 °C. Then, chromatin was pre-cleared with 30 µL Dynabeads Protein G (ThermoFisher, #10004D) by gentle end-to-end mixing at 4 °C for 2 h. Pre-cleared chromatin was diluted ten-fold in dilution buffer (50 mM Tris pH 8.0, 1 mM EDTA, 150 mM NaCl, 0.1% Triton X-100) and 5% (by volume) was saved as input until later. H3K27me3 antibody (4 µg, Millipore-Sigma, #07-449) was added to pre-cleared chromatin (30 µg) and mixed end-to-end at 4 °C overnight. The next day, antibody-chromatin complexes were captured by adding 50 µL Dynabeads Protein G and gentle end-to-end mixing at 4 °C for 2 h. They were washed once with low salt buffer (20 mM Tris pH 8.0, 2 mM EDTA, 150 mM NaCl, 1% Triton X-100, 0.1% SDS), once with high salt buffer (20 mM Tris pH 8.0, 2 mM EDTA, 500 mM NaCl, 1% Triton X-100, 0.1% SDS), once with LiCl wash buffer (10 mM Tris pH 8.0, 1 mM EDTA, 0.25 M LiCl, 1% NP-40, 1% sodium deoxycholate) and twice with TE buffer (10 mM Tris pH 8.0, 1 mM EDTA). For each wash, the immunoprecipitated complexes were mixed end-to-end at 4 °C for 5 min. Antibody-chromatin complexes were eluted from Dynabeads by incubating in elution buffer (0.1 M NaHCO_3_, 1% SDS) at 65 °C and vortexing. To the elution (ChIP) and input (saved earlier), NaCl was added to a final concentration of 200 mM to reverse crosslinked protein-DNA complexes and incubated overnight at 65 °C. The next day, the suspension was mixed with RNaseA and proteinase K to digest RNA and proteins, respectively, and incubated at 45 °C. Finally, ChIP and input chromatin were purified with Monarch PCR/DNA cleanup kit (NEB, #T1030) and eluted with H_2_O. The following protease inhibitors were added immediately before use to all of the above buffers: 250 µM Na3VO4, 1 mM NaF, 0.1 mM PMSF, 5 µg/mL aprotinin and leupeptin. DNA yield was quantified with Qubit fluorometer (ThermoFisher). Following the manufacturer’s recommendation, input (25 ng) or ChIP (0.5 ng) DNA were used to perform end repair, adaptor ligation and PCR amplification with NEBNext Ultra II DNA library prep kit and Multiplex Oligos (NEB, #E7645, #E7600). AmpureXP beads (Beckman Coulter, #A63880) were used at 0.9–1.0× reaction volumes for size selection. Input or ChIP samples were amplified by ten or fifteen PCR cycles, respectively, which yielded ∼600 ng DNA. Libraries were pooled and sequenced on NextSeq 500 at the London Regional Genomics Center with a high output 75 cycle kit to yield paired-end 38-bp read pairs.

### ChIP sequencing analysis

Demultiplexed FASTQ files were downloaded from BaseSpace. First, ENCODE blacklist regions ^79^ were filtered out from subsequent analysis. Reads were mapped to mm10 genome with bowtie2/2.4.2 with --sensitive-local setting. Samtools/1.12 was used with -f 0×2 option to keep concordantly mapped read pairs. Those read pairs were converted to sorted, indexed bam files. Biological duplicates were pooled together and MACS2/2.2.7.1 ^80^ was used to call broad peaks, outputting broadPeak files. A subcommand of deepTools/3.5.1 ^81^, bamCoverage, was used to generate reads-per-genomic-content (RPGC) normalized signal track (bw files) for all sequenced libraries. To account for composition biases in ChIP libraries, the trimmed mean of M-values method available in csaw/1.32.0 was used to generate scaling factors, which were passed into bamCoverage subcommand with --scaleFactor ^82, 83^. csaw was also used to quantify the loss of H3K27me3 at regions intersecting with genes and promoters upon GSK343 treatment compared to vehicle. plotEnrichment, computeMatrix, plotHeatmap and plotProfile commands (deepTools/3.5.1) were used to make barplots, heatmaps and profiles. bedtools/2.30.0 ^84^ was used to find peaks or number of peaks that intersected with RepeatMasker features, or were unique or common between treatment conditions or genotypes. ChIPseeker ^85^ was used to annotate peaks based on known genomic features. To show read mapping (colour-coded by strand) in Fig. S4B, samtools/1.12 was used with -f 0×40 option to split read pairs into two bam files. A bam file of one of the read mates (a ChIP sample of DMSO-treated WT B cells) was imported to IGV/2.11.1 ^86^ to show mapped reads at two loci. To show H3K27me3 signal enrichment at indicated loci (RPGC), bw files of biological replicates were merged together with bigWigMerge and bedGraphToBigWig ^87^. Merged signal track and broadPeak files were visualized on IGV/2.11.1. Peaks with a score less than 20 were filtered out.

### Flow cytometry

Total splenocytes or bone marrow cells were isolated as described above. 2 x 10^6^ cells were resuspended in PBS with ZombieNIR (1:500, BioLegend, #423105) and incubated on ice for 30 min to stain dead cells. Then, they were washed with FACS buffer and resuspended in pre-staining mix containing 50 µL Brilliant Stain Buffer (BD Biosciences, #563794), 5 µL TruStain monocyte blocker (BioLegend, #426102), 1 µL TruStain FcX plus (BioLegend, #156603) and 27.4 µL FACS buffer. Splenocytes were stained with the following antibody cocktail: anti-CD45.2 BV421 (BioLegend, #109831), anti-CD19 BV510 (BioLegend, #115545), anti-Ly6C BV605 (BioLegend, #128035), anti-CD11b FITC (BioLegend, #101206), anti-CD3ε PerCP/Cy5.5 (BioLegend, #100327), anti-CD8α PE (BioLegend, #100707) and anti-Ly6G APC (BioLegend, #127613). Bone marrow cells were stained with a similar panel but with the following substitution: anti-CD43 PerCP/Cy5.5 (BioLegend, #143219) and anti-CD45R PE (BioLegend, #103207). Stained cells were kept on ice for 30 min, then washed with FACS buffer. Cells were fixed in 100 µL fixation buffer (BD Biosciences, #554655) on ice for 20 min. They were washed and resuspended in 500 µL FACS buffer and 100 µL Precision counting beads (BioLegend, #424902). Flow cytometry was performed on LSR II (BD Biosciences) at the London Regional Flow Cytometry Facility and analyzed with FlowJo/10.8.1. AbC total antibody compensation beads and ArC amine reactive compensation beads (ThermoFisher, #A10497, #A10628) were used to generate compensation matrices in FACSDiva (BD Biosciences).

### Protein extraction and pulldown assay

Erythrocyte-lysed splenocytes were lysed in RIPA buffer (50 mM Tris pH 7.4, 150 mM NaCl, 1% NP-40, 0.5% sodium deoxycholate, 0.1% SDS, supplemented with protease inhibitors as above) on ice for 10 min. Chromatin fractions were prepared by lysing cells sequentially in buffer A (10 mM Tris pH 8.0, 10 mM KCl, 1.5 mM MgCl_2_, 0.34 M sucrose, 10% (v/v) glycerol, 0.1% Triton-X, supplemented with protease inhibitors as above), then in buffer B (3 mM EDTA, 0.2 mM EGTA, supplemented with protease inhibitors as above) with occasional mixing. Cytoplasmic and nucleoplasmic fractions were discarded by centrifugation, and resulting chromatin was digested with DNaseI in digestion buffer (20 mM Tris pH 7.5, 10 mM MgCl2).

For cGAS pulldown assay, erythrocyte-lysed splenocytes were lysed in non-ionic buffer (25 mM HEPES, 100 mM NaCl, 1 mM EDTA, 10% (v/v) glycerol, 1% Triton X-100, supplemented with protease inhibitors as above) on ice for 10 min. Debris was cleared by centrifugation at 16 000 x g for 30 min at 4 °C. Bradford assays were used to quantify protein concentration. For the pulldown assay, 500 µg of extract was mixed with 400 pmol of 5’ biotinylated ISD45 dsDNA and gently mixed end-to-end overnight at 4 °C. The next day, 20 µL streptavidin Dynabeads (ThermoFisher, #11205D) were added and gently mixed for 2 h at 4 °C. Pulldown complexes were washed once with non-ionic lysis buffer and once with non-ionic lysis buffer supplemented with 300 mM NaCl.

For histone acid extraction, purified, cultured splenic B cells were lysed in buffer (PBS, 0.5% Triton X-100, supplemented with protease inhibitors as above) on ice with gentle mixing for 10 min. Nuclei were isolated by centrifugation at 2000 RPM for 10 min at 4 °C. The pellet was resuspended in 0.4 N HCl overnight at 4 °C. Next day, cell debris was cleared by centrifugation, and trichloroacetic acid (1/3 volume) was added to the supernatant, and kept on ice for 2 hr. Precipitated histones were washed twice with 0.1% HCl (v/v) in cold acetone, then cold acetone. Residual acetone was removed by incubation at 55 °C. Histone pellet was resuspended in water before quantification with Bradford assays.

### SDS-PAGE and Western blot

RIPA extracts or pulldown samples were denatured by adding Laemmli buffer to 1x and boiling at 95 °C for 5 min. SDS-PAGE was performed following standard procedures. Samples were transferred to PVDF membrane using TransBlot Turbo (Biorad) and blocked in 5% skim milk in TBST for 1 h at RT. Membranes were incubated overnight at 4 °C with gentle shaking with the following primary antibodies: α-tubulin (CST, #2125, 1:5000), cGAS (CST, #31659, 1:1000), MDA5 (CST, #5321, 1:1000), RIG-I (Santa Cruz, #376845, 1:1000). The next day, membranes were washed five times with TBST and incubated with the following secondary antibodies for 1 h at RT with gentle shaking: m-IgG Fc BP-HRP (Santa Cruz, #525409, 1:500) or AffiniPure goat anti-rabbit IgG-HRP (JIR, #111-035-144, 1:5000-10000). After five washes with TBST, membranes were incubated in SuperSignal WestDura (ThermoFisher, #34075) and visualized on a ChemiDoc (Biorad). Coomassie staining was performed with GelCode Blue (ThermoFisher, #24590).

### CRISPR-Cas9 generation of RIC mutant mice

*In vitro* production of gRNAs and one-cell embryo injection were all performed as previously described ^56^. C57BL/6Crl mice were used as embryo donors, surrogate females and stud males. For restriction fragment length polymorphism assays, each of the three target loci was PCR amplified from tail DNA with Phire Animal Tissue Direct PCR kit (ThermoFisher, #F140WH). These products were either left undigested or restriction digested with the indicated enzymes and resolved on an agarose gel and EtBr stained. PCR reactions were also cloned into a plasmid and Sanger sequencing was performed to find the exact deletions in all mutant alleles present in the colony (Table S1). In addition, potential off-target sites were predicted *in silico* ^57^ and the two highest ranked intergenic and intragenic loci were genotyped through PCR amplification and sequencing (Table S2).

## Quantification and statistical analysis

All statistical analyses were performed with Prism 9. All relevant details are described in the figure legends.

## Supplemental item titles

Table S1 RIC mutant mice mutant alleles, related to Fig. 3

Table S2 RIC mutant mice off-target alleles, related to Fig. 3

Table S3 RNA-seq GSEA tables, related to Fig. 5.

Table S4 oligonucleotides

---

**Fig. S1.**
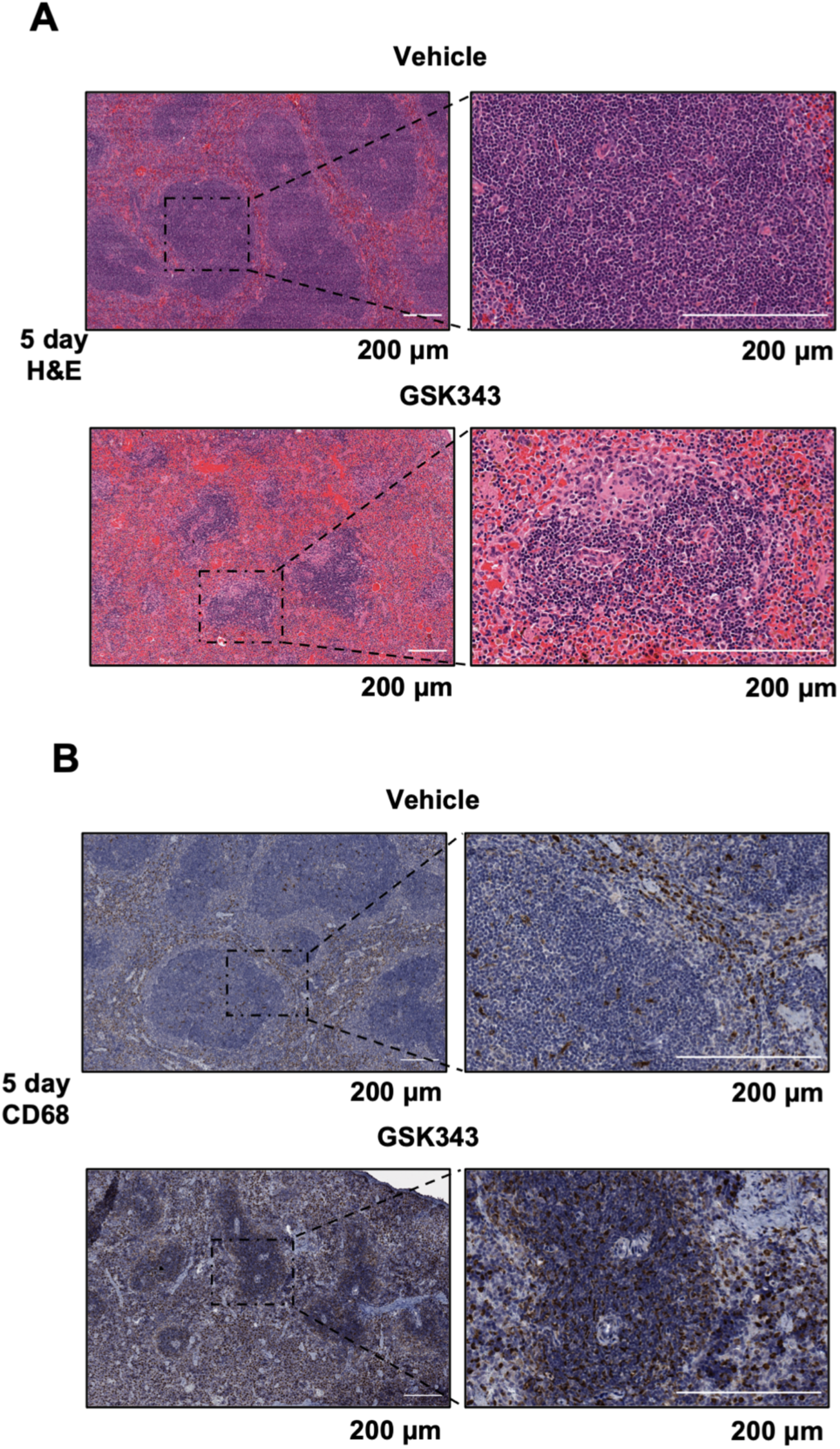
GSK343 induces inflammation and B cell death in the spleen. (A) H&E staining of spleens following 5 days of vehicle or GSK343 I.P injections. (B) CD68 staining of spleens from vehicle or GSK343 treated mice.

**Fig. S2.**
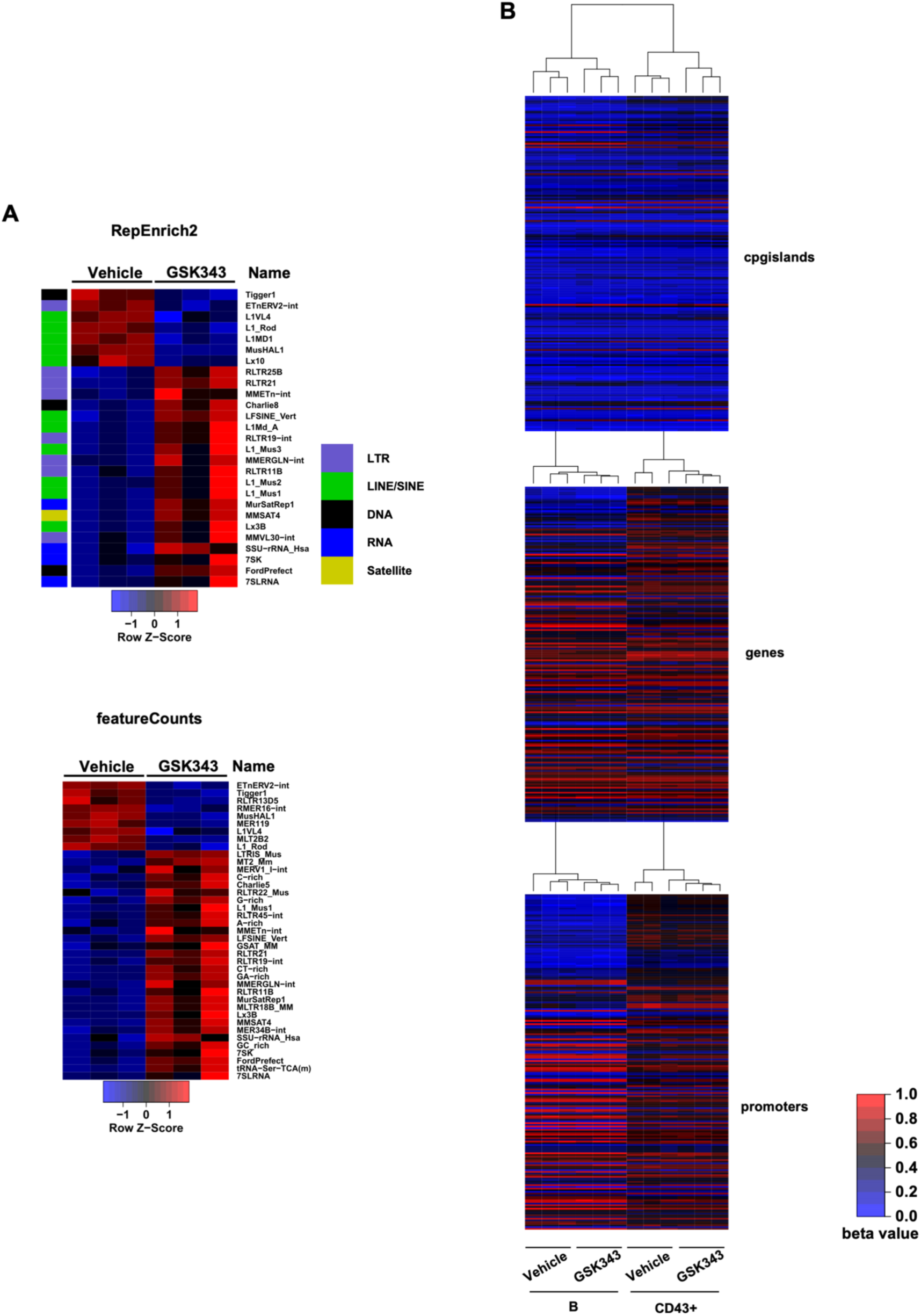
GSK343 upregulates repetitive element expression in splenic B cells. (A) Heatmaps of differential repetitive element expression in splenic B cells treated with either DMSO or 1 µM GSK343 for 48 h. Expression is shown as a Z-score of the mean of each row. RepEnrich2 pipeline is shown on the left, and featureCounts pipeline is shown in the right. (B) Heatmaps of DNA methylation probe beta values at top 200 differentially methylated probes annotated with CpG islands, genes or promoters between DMSO-treated B and CD43+ cells.

**Fig. S3.**
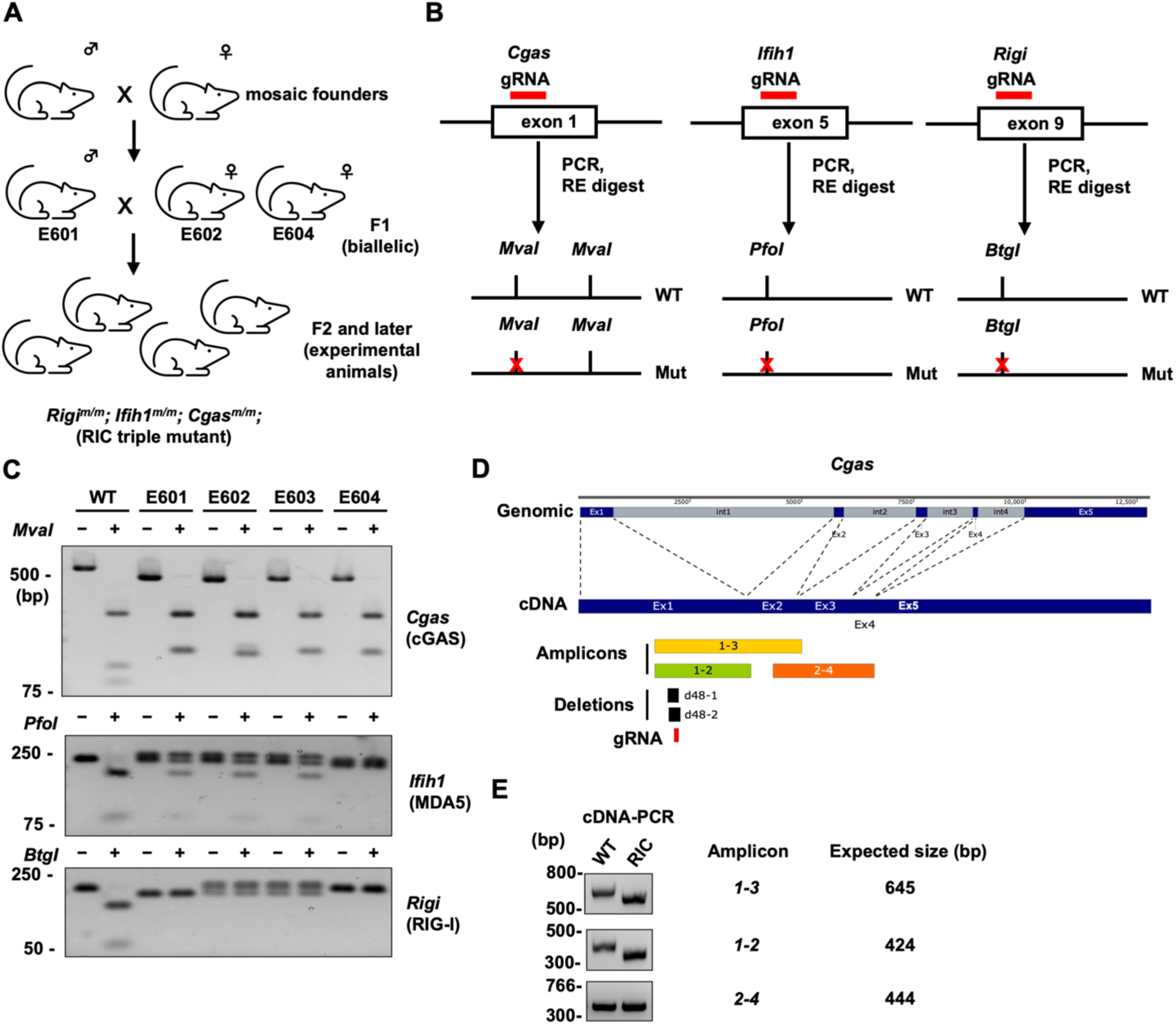
Strategy to generate and genotype RIC mutant mice. (A) Analysis of biallelic F1 mutant mice (E601, E602 and E604) that were used to generate experimental RIC mutant animals. (B) Restriction fragment length polymorphism (RFLP) assays to distinguish between WT and mutant alleles for each targeted gene. Upon CRISPR-Cas9 mediated DSB and non-homologous end joining, indicated restriction digest sites were disrupted. gRNA-targeted loci were PCR amplified and restriction digested. Resulting fragments were resolved by agarose gel electropheresis. (C) EtBr stained gels showing RFLP between WT and F1 mutants. Tail DNA was used to PCR amplify target exons of each gene, followed by digestion with the indicated restriction enzymes. (D) The structure of *Cgas* gene and mRNA, as well as PCR amplicons used to map deletions and gRNA recognition sites. (E) EtBr stained gels showing *Cgas* cDNA amplicons of WT and RIC mutant B cells as described in (D).

**Fig. S4.**
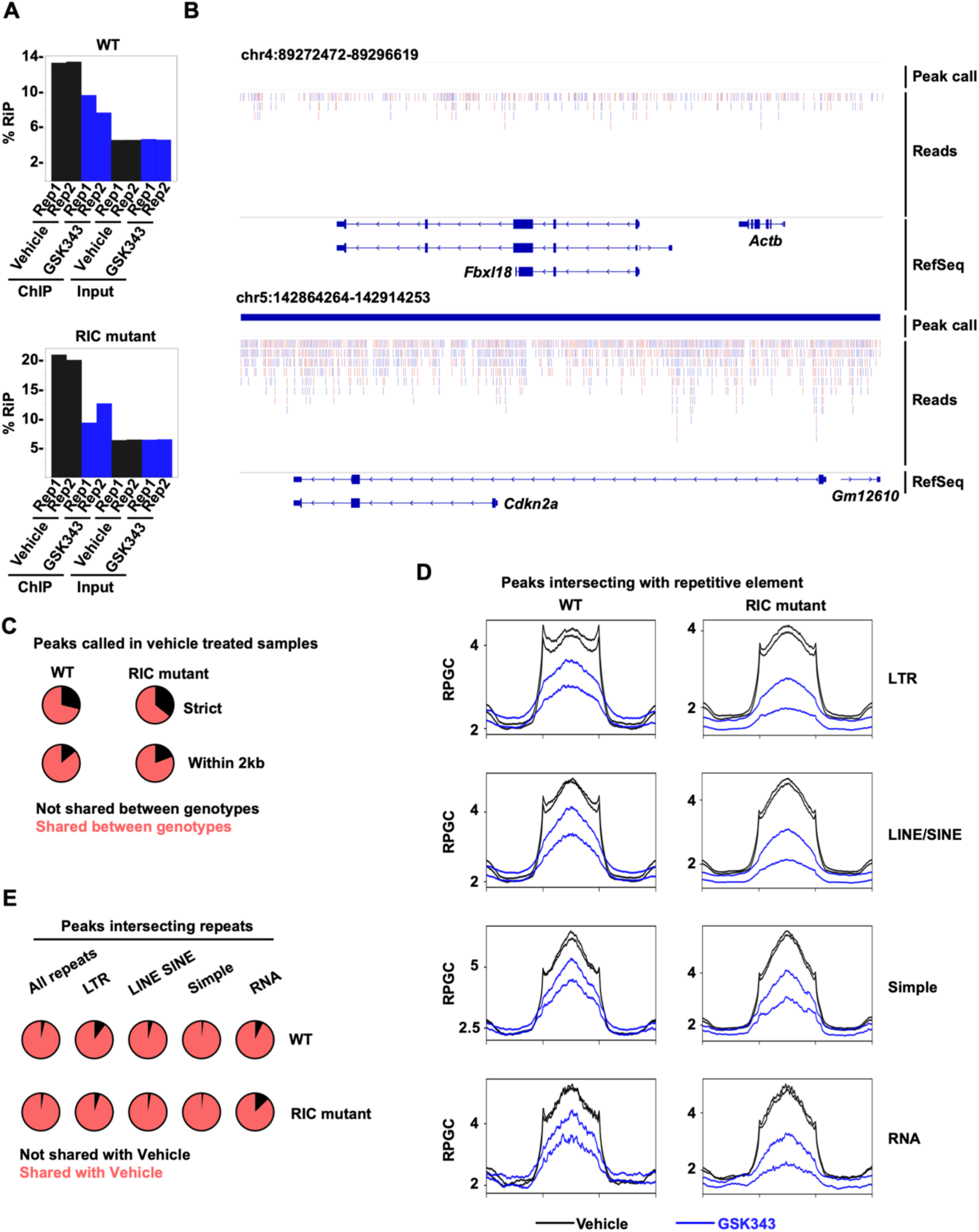
Similar GSK343 effects on H3K27me3 in WT and RIC mutants. (A) % reads in peaks (RiP) of all sequenced libraries. (B) Genome track view showing read alignment color-coded by alignment strand at one negative (top) and one positive (bottom) control locus. (C) Proportions of total peaks called in DMSO treated samples that are either not shared or shared between genotypes (top). Proportions of peaks called in DMSO treated samples that are either not shared or shared within 2 kb of each other between genotypes (bottom). (D) Enrichment profiles of H3K27me3 ChIP-seq reads at peaks called in DMSO or GSK343 treated samples intersecting with the indicated repetitive elements. Each box shows the average profile of scaled repetitive elements with 1 kb flanking each end. Biological duplicates for each treatment condition are shown as separate curves. (E) Proportions of total peaks intersecting with the indicated repetitive elements called in GSK343 treated samples that are either shared or not shared with DMSO treated samples.

**Fig. S5.**
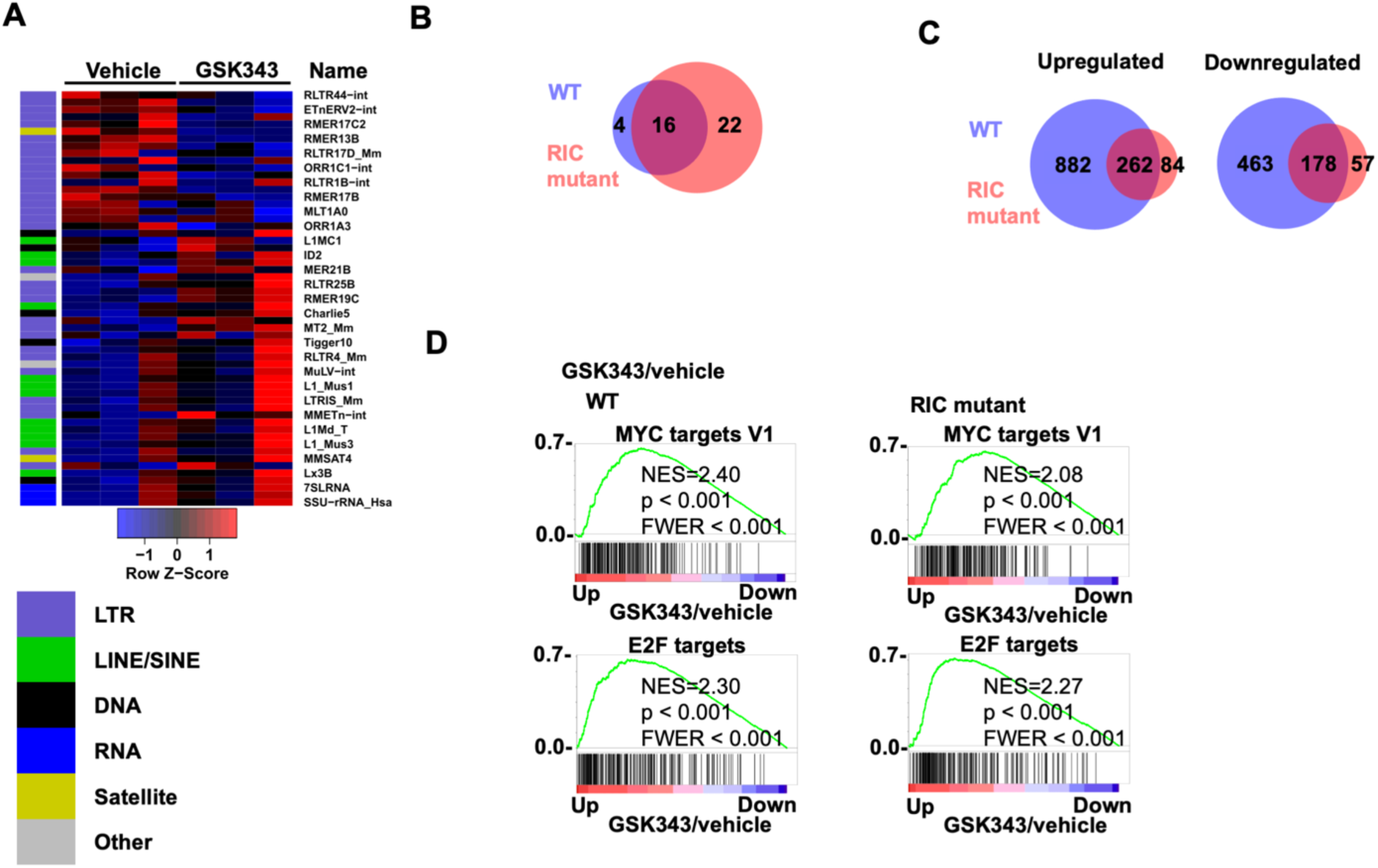
Expression profiles that are shared between WT and RIC mutant B cells upon GSK343 treatment. (A) Heatmap of differential repetitive element expression of three RIC mutant splenic B cell cultures treated with either DMSO or 1 µM GSK343 for 48 h. Expression is shown as a Z-score of the mean of each row. The leftmost column annotates each repeat into its repClass. (B) Venn diagrams showing the number of upregulated repetitive elements upon GSK343 treatment between WT and RIC mutant splenic B cells. (C) Venn diagrams showing the number of up or downregulated genes upon GSK343 treatment between WT and RIC mutant splenic B cells. (D) GSEA plots of Hallmark pathways comparing GSK343 to DMSO in WT and RIC mutant B cells.

